# Regulator of G protein signaling 6 (RGS6) in ventral tegmental area (VTA) dopamine neurons promotes EtOH seeking, behavioral reward and susceptibility to relapse

**DOI:** 10.1101/2023.10.24.563844

**Authors:** Mackenzie M. Spicer, Matthew A. Weber, Zili Luo, Jianqi Yang, Nandakumar S. Narayanan, Rory A. Fisher

## Abstract

Mesolimbic dopamine (DA) transmission is believed to play a critical role in mediating reward responses to drugs of abuse, including alcohol (EtOH). EtOH is the most abused substance worldwide with chronic consumption often leading to the development of dependence and abuse. Unfortunately, the neurobiological mechanisms underlying EtOH-seeking behavior and dependence are not fully understood, and abstinence remains the only effective way to prevent alcohol use disorders (AUDs). Here, we developed novel RGS6^fl/fl^; DAT-iCreER mice to determine the role of RGS6 in VTA DA neurons on EtOH consumption and reward behaviors. We found that RGS6 is expressed in DA neurons in both human and mouse VTA, and that RGS6 loss in mice upregulates DA transporter (DAT) expression in VTA DA neuron synaptic terminals. Remarkably, loss of RGS6 in VTA DA neurons significantly reduced EtOH consumption, preference, and reward in a manner indistinguishable from that seen in RGS6^−/−^ mice. Strikingly, RGS6 loss from VTA DA neurons before or after EtOH behavioral reward is established significantly reduced (∼50%) re-instatement of reward following extinguishment, demonstrating distinct roles of RGS6 in promoting reward and relapse susceptibility to EtOH. These studies illuminate a critical role of RGS6 in the mesolimbic circuit in promoting EtOH seeking, reward, and reinstatement. We propose that RGS6 functions to promote DA transmission through its function as a negative modulator of GPCR-Gα_i/o_-DAT signaling in VTA DA neurons. These studies identify RGS6 as a potential therapeutic target for behavioral reward and relapse to EtOH.

## 1. Introduction

Alcohol (EtOH) is the most commonly abused drug worldwide and alcohol use disorders (AUDs) impart a significant socioeconomic burden on the U.S. healthcare system, affecting ∼12% of the population. The neurobiological mechanisms underlying alcohol-seeking behavior and dependence are not fully understood, resulting in limited therapeutics. While there are currently several approved therapeutics designed to reduce alcohol cravings or withdrawal symptoms, abstinence remains the only effective way to prevent AUDs.

Chronic EtOH exposure causes an amalgamation of neuroadaptations resulting in the development of alcohol dependence. EtOH functions as a CNS depressant to simultaneously dampen NMDA and enhance GABA_A_R function(1). Moreover, accompanying changes in glutamatergic and GABAergic inputs promote acute neurotransmitter abnormalities and chronic alterations in synaptic plasticity within the mesolimbic circuit(2–8). The mesolimbic circuit is comprised of dopamine (DA) neurons within the ventral tegmental area (VTA) projecting to GABAergic neurons in the nucleus accumbens (NAc) and is a major dopaminergic signaling pathway heavily implicated in drug abuse and addiction biology. Numerous studies have shown that EtOH self-administration in rodents promotes VTA dopamine (DA) release in the NAc, and that pharmacological manipulations of DA actions on D_1_ and D_2_ medium spiny neurons (MSNs) of the NAc (which account for more than 90% of the neurons in the NAc) alter responsiveness to EtOH(2, 3, 5, 6, 9–11). Furthermore, EtOH self-administration directly to the VTA is reduced by stimulation of dopaminergic autoreceptors (D_2_Rs) in the VTA(12). This latter finding highlights the central role of G protein-coupled receptor (GPCR) signaling in presynaptic VTA DA neurons in modulating the activity of the mesolimbic circuit and EtOH consumption.

Several neurotransmitters (DA; GABA; opioids and serotonin) modulate neuronal communication in the mesolimbic pathway by activating a variety of GPCRs (D_2_Rs, GABA_B_Rs, μ-opioid receptors (MORs), and serotonin 1A receptors (5-HT_1A_Rs)). Abnormalities in these GPCRs have been implicated in alcohol dependence in human AUD(13–17). Drugs targeting the GABA, opioid, DA, and serotonin neurotransmitter systems have been evaluated for clinical utility in the treatment of alcoholism(18–20). Of the three FDA-approved drugs for AUD, naltrexone (an opioid receptor antagonist) is the only drug that acts upon the mesolimbic pathway to limit the rewarding actions of EtOH. Naltrexone blocks opioid inhibition of GABAergic interneurons in the VTA, thereby promoting their inhibition of VTA DA neurons(21). Thus, elucidating the GPCR-regulated mechanisms underlying alcohol dependence may lead to the development of novel therapies for AUD. Here, we investigate the action of a critical modulator of G protein signaling in VTA DA neurons in alcohol behaviors.

Regulators of G protein Signaling (RGS) proteins function as essential negative modulators of GPCR signaling(22–25). The semiconserved RGS domain of RGS proteins confers functional GTPase-activating protein (GAP) activity towards specific Gα subunits, thereby accelerating the turn-off mechanism for G protein signaling and reducing the magnitude and duration of G protein signaling. RGS6, a member of the R7 RGS protein family originally cloned in our laboratory(26) (Genbank AF073920, 1988), exhibits GAP activity specifically toward Gα_i/o_(27) G proteins, which are activated by numerous GPCRs in the mesolimbic circuit implicated in human AUDs. We previously identified RGS6 as a critical negative regulator of Gα_i/o_-coupled GABA_B_Rs, 5-HT_1A_Rs and D_2_Rs(28–30), and RGS6 has been shown to also function as a regulator of MOR signaling(31).

We previously discovered that RGS6 promotes EtOH-seeking and reward behaviors through its ability to negatively regulate neuronal Gα_i/o_-coupled GPCRs implicated in AUDs(32, 33). We found that RGS6^−/−^ mice consume less EtOH when given free access, and are less susceptible to EtOH reward, dependence, and withdrawal. Additionally, antagonism of GABA_B_Rs or D_2_Rs partially reversed the reduced EtOH consumption observed in RGS6^−/−^ mice, whereas DAT inhibition completely restored their EtOH consumption. EtOH consumption also specifically induced the expression of RGS6 in the VTA, a potential mechanism underlying chronic EtOH consumption/dependence. Furthermore, RGS6 deficiency was associated with reduced striatal DA levels, suggesting that RGS6 may regulate DA signaling and/or availability. A year following our report, a GWAS identified *RGS6* as one of four novel genes linked to human AUDs(33). Taken together, these findings implicate RGS6 as an essential regulator of EtOH consumption and reward, DA bioavailability, and AUDs.

Here, we sought to determine the role of RGS6 expression in VTA DA neurons on EtOH behaviors and mesolimbic DA signaling. We discovered that RGS6 is expressed in the VTA DA neurons of both humans and mice. Using a tamoxifen-inducible Cre mouse model to allow for DAT-specific neuronal loss of RGS6, we discovered that VTA DA neuron RGS6 expression critically regulates free EtOH consumption, preference, reward, and reinstatement. Importantly, RGS6 deletion after establishing EtOH reward prevented reinstatement-induced relapse. Moreover, RGS6 co-localizes with DAT in VTA DA neuron cell bodies and synaptic terminals in the NAc, and RGS6 loss increases DAT expression at both locations. These results demonstrate that RGS6 functions as an essential modulator of Gα_i/o_ signaling in VTA neurons and through this action modulates both EtOH seeking/preference, reward, and the transition to dependence/addiction.

### 2. Methods

### 2.1 Mice

All behavioral studies employed RGS6 floxed (RGS6^fl/fl^); DAT-iCreER mice treated with sunflower oil (control; RGS6 WT^VTA^) or tamoxifen (75 mg/kg, i.p. daily for 5 days; RGS6 KO^VTA^), with RGS6^+/+^ (C57BL6/J) and global RGS6^−/−^ mice (C57BL6/J background) serving as controls. RGS6^fl/fl^ mice on a mixed background (C57BL/6J x SJL) were created using CRISPR-Cas9 gene editing to target exon 5 for deletion of all known RGS6 splice forms(26, 34). RGS6^fl/fl^ mice were bred with DAT-iCreER mice (Jackson Laboratories #016583) to create mice to determine the impact of RGS6 loss in VTA DA neurons on EtOH seeking and reward behaviors, and DA neuron activity. All *in vivo* experiments were approved and performed in accordance with guidelines set by the Institutional Animal Care and Use Committee (IACUC) at the University of Iowa. Protocols describing behavioral studies, histology, and immunoblotting can be found in “*Supplementary Materials*.”

### 2.5 Statistical Analysis

All data are expressed as mean ± SEM. Student’s T test was used to analyze IF and WB quantification data (**Fig. 1**; **Fig. 4; Suppl. Fig. 1**). Two-way ANOVA with Tukey’s *post-hoc* adjustment was used to analyze the effects of and interaction between genotype and sex (**Fig. 2**; **Fig. 3**). Multi-way ANOVA with Tukey’s post-hoc adjustment was used to analyze the effects of and interaction between genotype, sex, and treatment (saline vs. EtOH) during CPP, extinguishment and reinstatement behavior assays (**Fig. 2**; **Fig. 3; Suppl. Fig. 2**). Sex as a biological variable was accounted for in all statistical analyses. Animal cohort sizes were determined from preliminary studies and G*Power analyses. *P* ≤ 0.05 was considered statistically significant. Statistical analyses were performed using XLSTAT software, and graphs were created using GraphPad Prism.

**Figure 1.**
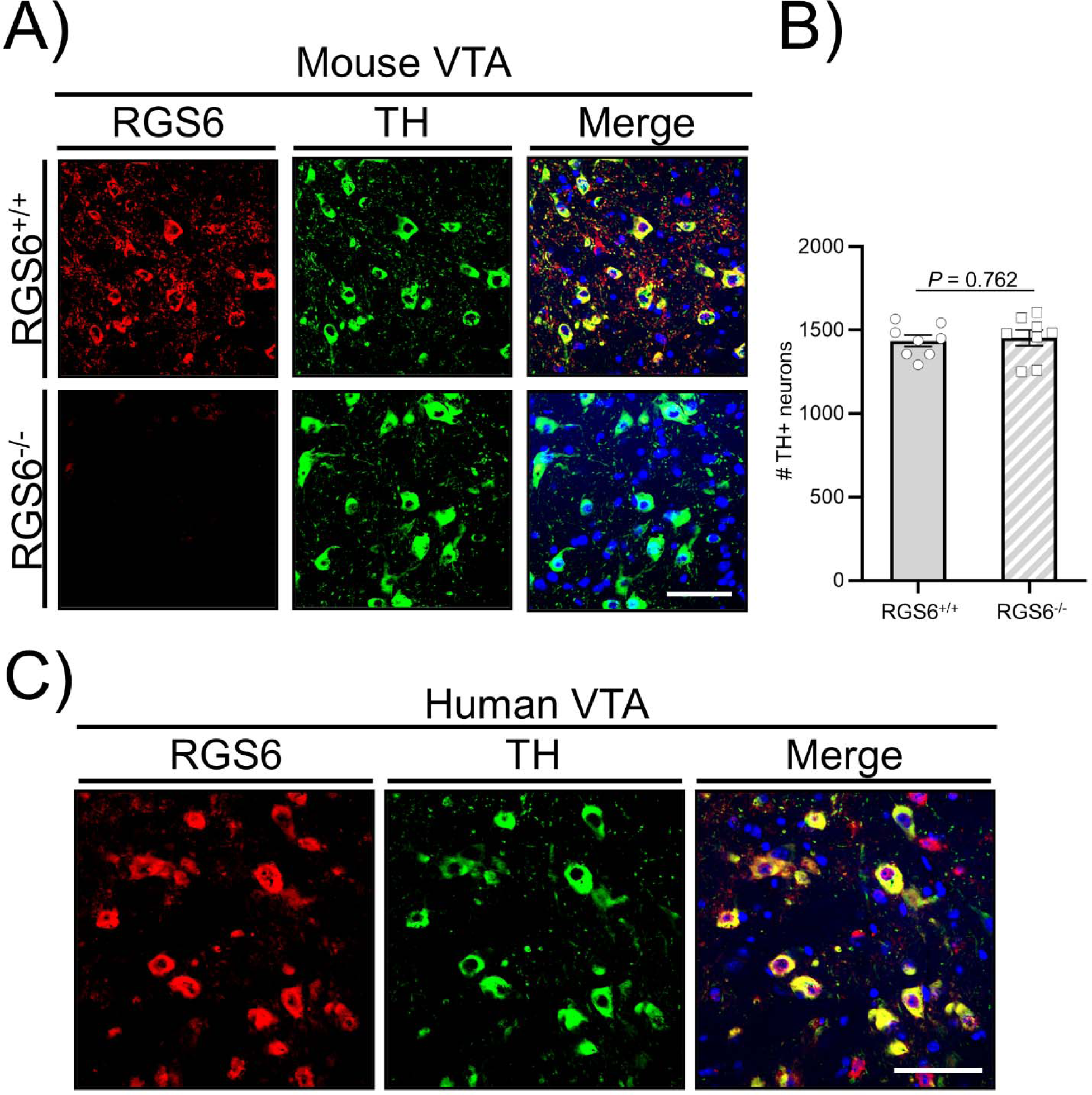
RGS6 is expressed in both human and mouse VTA dopamine neurons. **A-B)** Immunofluorescent (IF) analysis (**A**) and quantification (**B**) of RGS6 (red) expression in dopamine neurons (TH+, green) in mouse VTA. **C)** IF analysis of RGS6 (red) expression in dopamine neurons (TH+, green) in human VTA. VTA = ventral tegmental area. Scale bars represent 40 µm.

**Figure 2.**
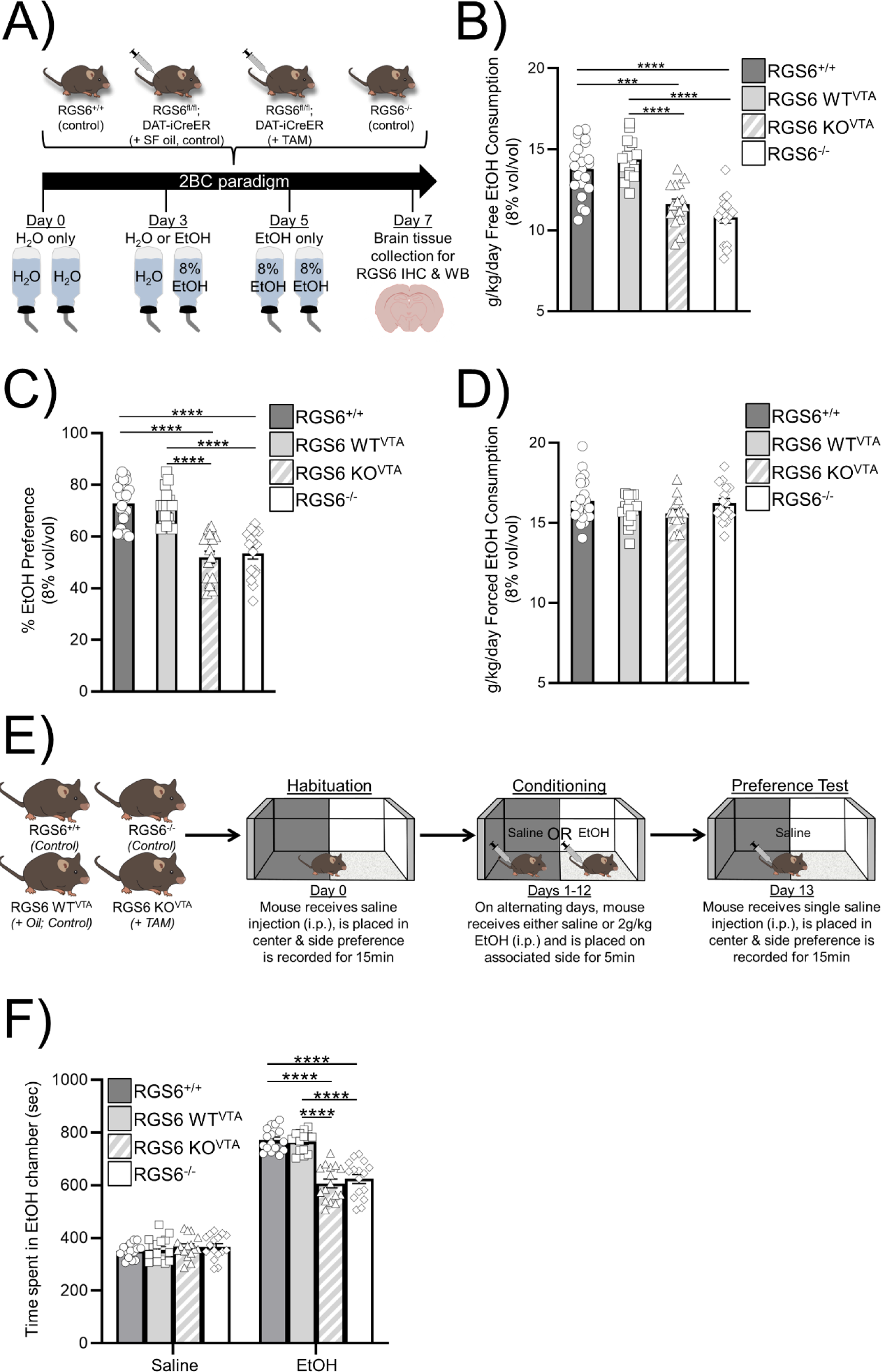
RGS6 expression in VTA DA neurons promotes EtOH drinking and reward behaviors. **A)** Schematic outlining experimental design of 2 bottle choice (2BC) acute EtOH consumption paradigm. **B-D)** Free EtOH consumption (**B**), EtOH preference (**C**), and forced EtOH consumption (**D**) of RGS6^+/+^, RGS6 WT^VTA^, RGS6 KO^VTA^, and RGS6^−/−^ mice during 2BC. **E)** Schematic depicting CPP chamber set up and experimental timeline. **F)** Recorded time spent in the EtOH chamber during the preference testing phase of CPP. Data are expressed as mean ± SEM. Two-way ANOVA with Tukey’s *post-hoc* adjustment was used to analyze EtOH consumption and preference data to examine the effects of and interactions between genotype and sex. A significant effect of genotype was observed for free EtOH consumption (*F*_3,61_ = 22.948, *P* = 0.000) and preference (*F*_3,61_ = 28.384, *P* = 0.000), but not forced EtOH consumption. No significant effects of sex or its interaction with genotype were observed for free EtOH consumption, preference, or forced EtOH consumption. Multiway ANOVA with Tukey’s *post-hoc* adjustment was used to analyze the effects of and interactions between genotype, treatment, and sex for CPP. Significant effects of and interactions between treatment and genotype were observed for CPP (Genotype: *F*_3,112_ = 23.538, *P* = 0.000; Treatment: *F*_1,112_ = 1575.568, *P* = 0.000; Genotype*Treatment: *F*_3,112_ = 33.164, *P* = 0.000). No significant effect of sex or its interactions were observed across all behaviors. \**P* ≤ 0.05, \*\**P* ≤ 0.01, \*\*\**P* ≤ 0.001, \*\*\*\**P* ≤ 0.0001.

**Figure 3.**
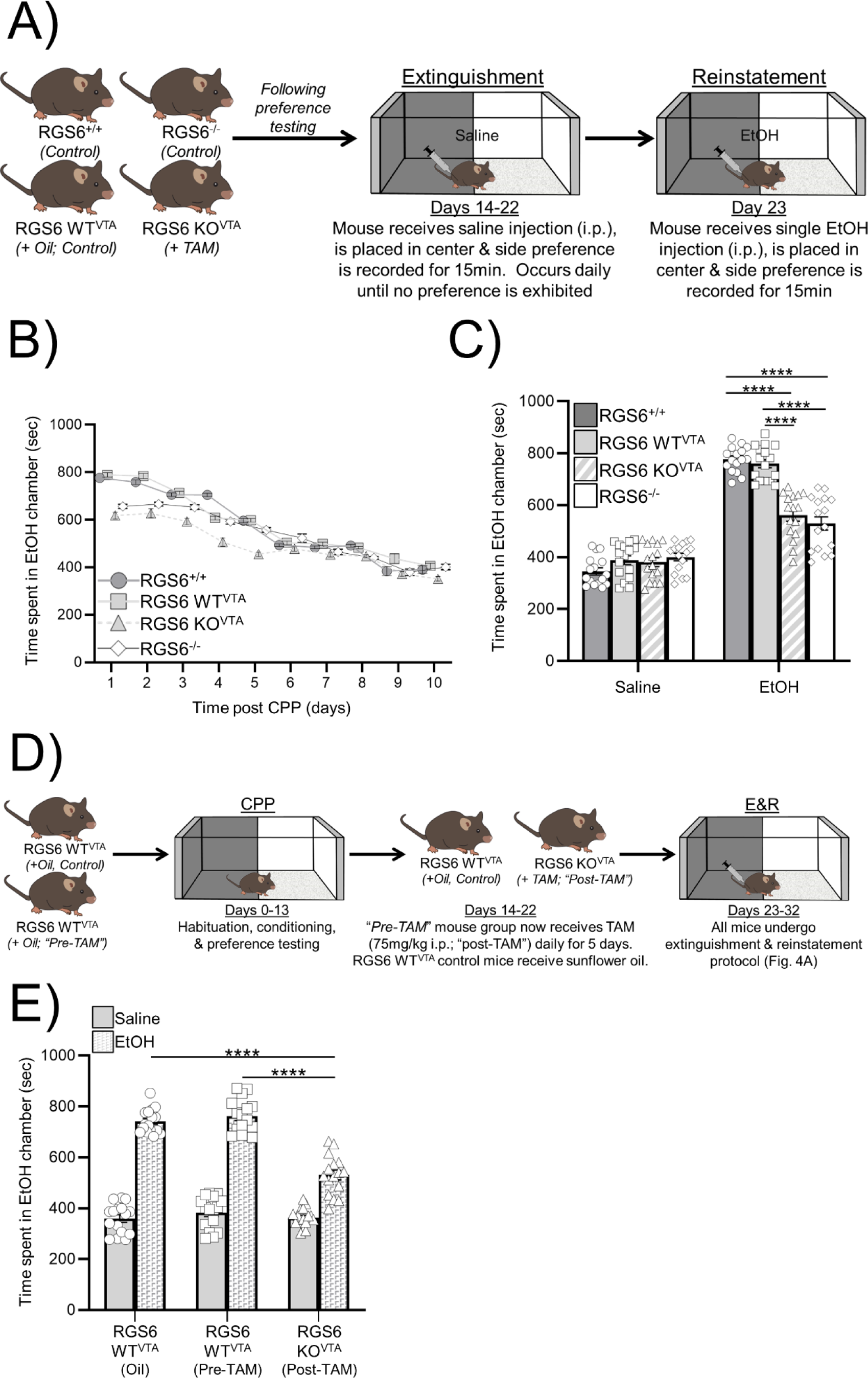
Distinct roles of RGS6 in VTA DA neurons on EtOH reward and relapse susceptibility. **A)** Schematic of extinguishment-reinstatement (E&R) paradigm for EtOH relapse following CPP-measured reward. **B)** Average daily recorded time spent in the EtOH chamber during extinguishment following CPP of mice from Fig. 3E. **C)** Time spent in the EtOH chamber during reinstatement following singular injection of EtOH (2 g/kg, i.p.) in these mice in which RGS6 deletion occurred before established CPP reward. **D-E)** Schematic of extinguishment-reinstatement paradigm for EtOH relapse with deletion of RGS6 following established CPP award (**D**) and time spent in the EtOH chamber during reinstatement following EtOH (2 g/kg, i.p.) in these mice (**E**). Data are expressed as mean ± SEM. Two-way ANOVA with Tukey’s *post-hoc* adjustment was used to examine the effects of and interactions between genotype and sex and reinstatement ((**B**): Genotype: *F*_3,112_ = 20.438, *P* = 0.000; Treatment: *F*_1,112_ = 470.821, *P* = 0.000; Genotype*Treatment: *F*_3,112_ = 32.536, *P* = 0.000; (**D**): Genotype: *F*_3,112_ = 39.107, *P* = 0.000; Treatment: *F*_1,112_ = 621.858, *P* = 0.000; Genotype*Treatment: *F*_3,112_ = 32.680, *P* = 0.000). No significant effect of sex or its interactions were observed. \**P* ≤ 0.05, \*\**P* ≤ 0.01, \*\*\**P* ≤ 0.001, \*\*\*\**P* ≤ 0.0001.

**Figure 4.**
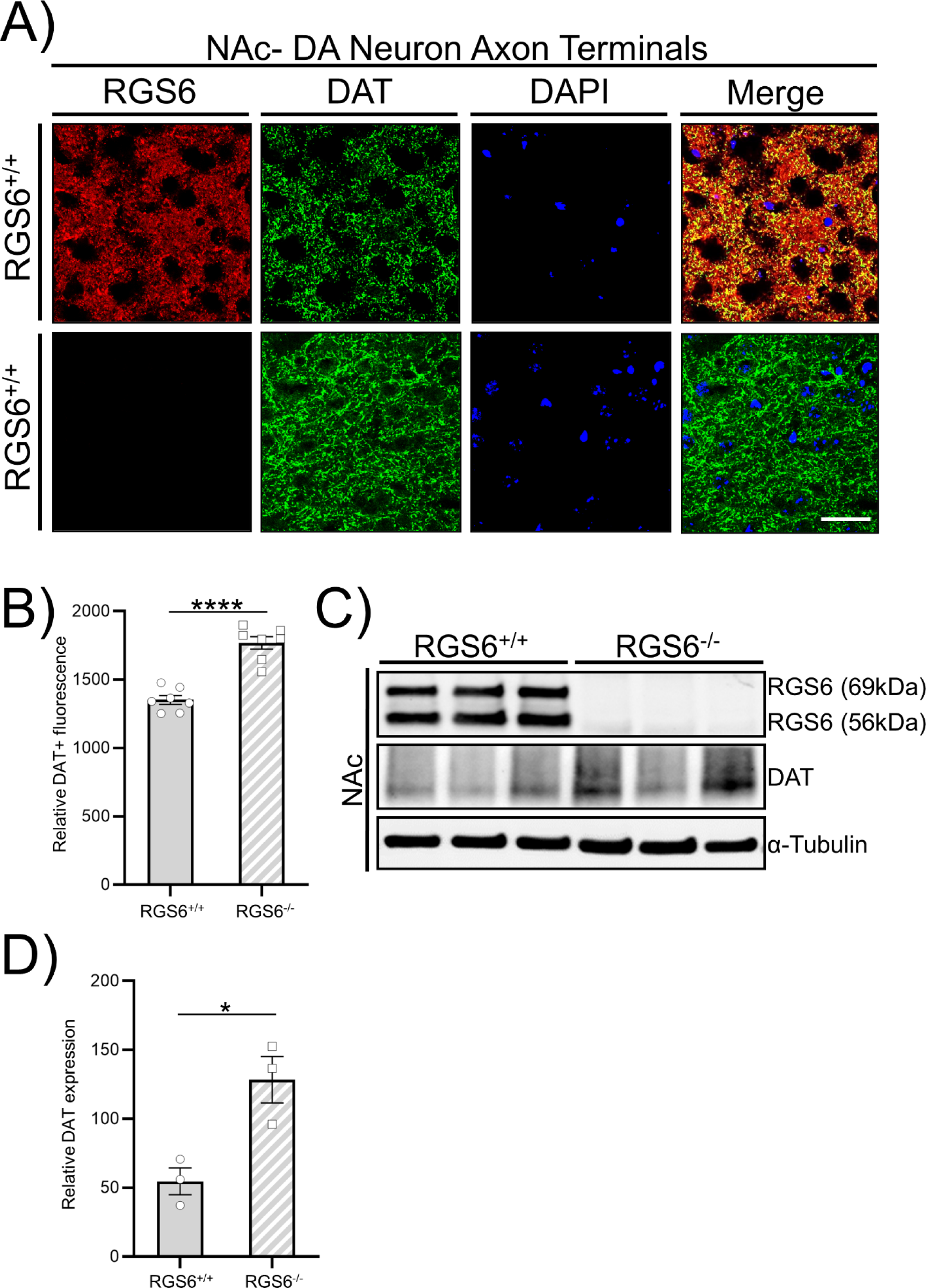
RGS6 loss leads to DAT upregulation in the VTA and NAc of mice. **A-B)** IF analysis of RGS6 (red) co-expression with DAT (green) in VTA DA neuron cell bodies (**A**) and synaptic terminals in the NAc (**B**) of 3mo. RGS6^+/+^ and RGS6^−/−^ mice. **C-D)** Western blot (WB) analysis (**C**) and quantification (**D**) of RGS6 expression on DAT expression in the NAc of 3mo. RGS6^+/+^ and RGS6^−/−^ mice (α-Tubulin = loading control). \**P* ≤ 0.05, \*\*\*\**P* ≤ 0.0001

## 3. Results

### 3.1 RGS6 is expressed in VTA DA neurons in mice and humans

EtOH’s hedonic value is driven by increased DA release from the VTA to the NAc, caused by EtOH’s ability to hijack the brain’s reward system(2, 6–8). Our previous study showed that mice with global deletion of RGS6 exhibit significantly reduced EtOH consumption, behavioral reward, dependence, and withdrawal to EtOH, as well as reduced DA bioavailability(32). In that study, we identified a population of RGS6-expressing DA neurons in mouse VTA. However, it remains unclear whether RGS6 is expressed in all DA neurons in mouse VTA or if it is similarly expressed in human VTA DA neurons, where *RGS6* was identified as an AUD gene(33) following our studies in mice. Therefore, we examined the expression of RGS6 in mouse and human VTA and developed mice to selectively delete RGS6 from VTA DA neurons to establish its role in EtOH seeking/consumption, reward, and addictive behaviors.

Immunofluorescent (IF) analysis revealed that RGS6 is expressed in all tyrosine hydroxylase (TH) positive DA neurons in the VTA of mice and humans. **Fig. 1A** compares the expression of RGS6 and TH in the VTA of wild-type (RGS6^+/+^*, upper*) and RGS6^−/−^ (*lower*) mice. As shown, RGS6 expression is found in TH^+^ neurons with the specificity of RGS6 staining shown by loss of RGS6 expression in these neurons in RGS6^−/−^ mice. We previously described an age-dependent loss of DA neurons beginning around 6mos in the substantia nigra pars compacta (SNc) but not the VTA(28, 35) in RGS6^−/−^ mice. In agreement with those studies, we did not observe loss of DA neurons in the VTA of 3mo. old RGS6^−/−^ mice (**Fig. 1B**). Therefore, the diminished EtOH consumption and reward behaviors in these mice(32) are not due to neurodegeneration. **Fig. 1C** shows that RGS6 is expressed in all TH^+^ neurons in the human VTA. Thus, both human and mouse VTA DA neurons express RGS6.

### 3.2 Selective deletion of RGS6 from VTA DA neurons

To determine the role of RGS6 in VTA DA neurons, we employed novel RGS6 floxed (RGS6^fl/fl^) mice in which we flanked exon 5 of the mouse *Rgs6* gene with loxP sites ((34). We targeted exon 5 because our analysis of the *Rgs6* gene determined that Cre-mediated deletion of this exon would promote loss of all 36 RGS6 isoforms identified in our initial cloning efforts(26). RGS6^fl/fl^ mice received bilateral injections of either AAV-Cre-eGFP or AAV-eGFP and were euthanized 3wks later to examine RGS6 expression in VTA by western analysis. **Supplemental Fig. 1A** shows that Cre-mediated deletion of RGS6 in these mice was successful, as AAV-Cre-eGFP treated RGS6^fl/fl^ mice showed loss of RGS6 expression in the VTA as observed in RGS6^−/−^ mice.

VTA DA neurons have been targeted using mouse lines expressing TH-Cre or DAT-Cre, however, Lammel *et al*.(36) demonstrated that DAT-Cre mice show specific targeting of VTA DA neurons while many midbrain neurons targeted by TH-Cre or TH-GFP are not TH positive. Also given that VTA neurons projecting to the NAc have high DAT levels compared to those projecting to non-striatal sites, DAT-Cre lines are the best choice for mesolimbic manipulation of RGS6, which is expressed exclusively in DA neurons in the VTA. Therefore, we then crossed these RGS6^fl/fl^ mice to a tamoxifen (TAM)-inducible dopamine transporter (DAT)-iCreER mouse model, creating RGS6^fl/fl^; DAT-iCreER mice. 3mo old DAT-iCreER, RGS6^fl/fl^ mice were then injected with tamoxifen or sunflower oil (control) and euthanized 3wks later to examine RGS6 expression in VTA DA neurons by IF. **Supplemental Fig. 1B** shows that, while RGS6 co-localized with DAT^+^ DA neurons in the VTA of control mice, tamoxifen treated mice showed complete loss of RGS6 from these neurons. Further, RGS6 deletion from VTA DA neurons did not alter the number of DA neurons (**Suppl. Fig. 1C**) in the VTA, consistent with our observation in RGS6^−/−^ mice (**Fig. 1B**).

### 3.3. RGS6 in VTA DA neurons drives EtOH consumption and reward/relapse behaviors

We performed studies to evaluate the effects of RGS6 deletion from VTA DA neurons on EtOH consumption and reward/dependence behaviors in mice. Behaviors in these mice (RGS6 KO^VTA^) were compared to those of WT (RGS6^+/+^), global KO (RGS6^−/−^), and control RGS6^fl/fl^ mice treated with sunflower oil (RGS6 WT^VTA^) mice. First, mice were subjected to the 2BC EtOH drinking paradigm (**Fig. 2A**) to examine EtOH consumption and preference. RGS6 KO^VTA^ mice, like RGS6^−/−^ mice, exhibited significantly reduced EtOH consumption and EtOH preference compared to RGS6^+/+^ and RGS6 WT^VTA^ mice (**Figs. 2B,C**). In contrast, no differences were observed in forced EtOH consumption in these groups of mice (**Fig. 2D**). Because the behavioral rewarding effects of EtOH are known to correlate with voluntary EtOH consumption and preference(37), we next examined EtOH-mediated conditioned reward in mice using conditioned place preference (CPP) behavioral assays (**Fig. 2E**). EtOH-induced CPP has been shown to be expressed through a VTA-dependent mechanism(38). Consistent with the reduction in EtOH consumption and preference, RGS6 KO^VTA^ mice showed reduced susceptibility to EtOH conditioned reward as seen in RGS6^−/−^ mice (**Fig. 2F**).

AUD is a chronic disorder with a high rate of relapse, with craving and relapse triggered by environmental cues(39–41). CPP induced by various drugs, including EtOH, can be extinguished and reinstated by drug priming(42). Given our findings that RGS6^−/−^ mice are less susceptible to EtOH dependence and withdrawal(32) and that RGS6^−/−^ and RGS6 KO^VTA^ mice show reduced CPP to EtOH (**Fig. 2E**), we performed experiments to examine effects of RGS6 loss on extinguishment and re-instatement of EtOH induced CPP in mice. We used an extinguishment-reinstatement paradigm, which models susceptibility to drug-induced reward and relapse(42), following CPP-measured reward (**Fig. 3A**). **Fig. 3B** shows the time spent in the EtOH chamber during the extinguishment paradigm by mice of different genotypes. As shown, all mice groups exhibited a time-dependent decline in the time spent in the EtOH chamber following CPP, with the reduced CPP in RGS6^−/−^ and RGS6 KO^VTA^ mice compared to control mice disappearing by day 6. We then performed re-instatement in these mice by singular injection of EtOH (2g/kg, i.p.). **Fig. 3C** shows that reinstatement of CPP following extinguishment was significantly and equivalently impaired in RGS6 KO^VTA^ and RGS6^−/−^ mice. We next asked whether deletion of RGS6 following CPP-established EtOH reward would interfere with reinstatement-induced relapse. To accomplish this, after completion of CPP and extinguishment, RGS6^fl/fl^; DAT-iCreER mice were divided into 2 subgroups: one receiving sunflower oil (control) and one that would eventually receive tamoxifen (**Fig. 3D**). In the latter group, reinstatement behavior was measured before (pre-TAM; control) and after (post-TAM) tamoxifen injection to assess the impact of RGS6 loss. Remarkably, RGS6 deletion from VTA DA neurons in mice following established reward significantly impaired reinstatement-induced relapse, as indicated by their decreased time spent in the EtOH-associated box (**Fig. 3E**, post-TAM) relative to their pre-TAM and oil controls.

Female C57BL/6 mice have been reported to have increased EtOH consumption compared to males. This has been observed in 2BC using 10% EtOH but not 3% EtOH(43) and in binge drinking paradigms or during long term EtOH drinking (>1wk) using high EtOH concentrations (>10%)(44–46). However, these female mice do not show enhanced behavioral reward to EtOH measured by CPP(47, 48). Though we observed no sex-specific effects on EtOH behaviors in WT and RGS6^−/−^ mice(32), we stratified our current EtOH behavioral results by sex to determine whether the effects of RGS6 VTA DA neuronal loss on these behaviors were sex-dependent. This analysis revealed that the reduced EtOH consumption in 2BC (8% EtOH) (**Suppl. Fig. 2A**), EtOH preference (**Suppl. Fig. 2B**), CPP (**Suppl. Fig. 2D**), and extinguishment (**Suppl. Fig. 2E**) and reinstatement of EtOH reward (**Suppl. Figs. 2F-G**) of RGS6 KO^VTA^ mice did not differ between sexes. Additionally, no differences were observed in forced EtOH consumption in RGS6 KO^VTA^ mice in either sex (**Suppl. Fig. 2C**).

These studies demonstrate that RGS6 loss from VTA DA neurons leads to a significant reduction in EtOH voluntary consumption, preference, behavioral reward, and susceptibility to relapse in both male and female mice.

### 3.4 RGS6 is expressed in VTA dopaminergic terminals and suppresses DAT upregulation, likely by serving as a negative regulator of the GPCR-Gα_i/o_-DAT signaling axis

Presynaptic Gα_i/o_-coupled D_2_ autoreceptors on VTA DA neurons block vesicular DA release and upregulate DAT expression through multiple mechanisms(49–53). The feedback of synaptic DA onto these receptors functions to reduce DA release and to increase DA clearance and resultant behavioral responses to DA in the mesolimbic circuit. Our previous study showed that diminished EtOH consumption of RGS6^−/−^ mice was partly reversed by antagonism of Gα_i/o_-coupled D_2_Rs or GABA_B_Rs while singular inhibition of DAT completely restored EtOH consumption in these mice(32).

These findings suggest that RGS6 suppression of DAT upregulation by presynaptic Gα_i/o_-coupled GPCRs is the primary mechanism by which RGS6 promotes EtOH seeking and reward behaviors. RGS6^−/−^ and RGS6 KO^VTA^ mice phenocopy each other in terms of exhibiting reduced EtOH consumption, preference, reward, and relapse behavior compared to RGS6^+/+^ mice (**Fig. 3; Suppl. Fig. 2**). Therefore, we compared DAT expression at synaptic terminals in the NAc by western blot and IF in RGS6^+/+^ and RGS6^−/−^ mice. **Fig. 4A** (upper) shows that RGS6 is co-expressed with DAT at VTA synaptic terminals in the NAc of RGS6^+/+^ mice. EtOH and other drugs of abuse induce DA release from these terminals which literally bathe the medium spiny neurons in the NAc. DAT expression in VTA synaptic terminals was significantly higher in RGS6^−/−^ mice compared to RGS6^+/+^ mice (**Fig. 4A** lower, **Fig. 4B**). This was confirmed by western analysis of DAT expression in the NAc of RGS6^+/+^ and RGS6^−/−^ mice (**Figs. 4C-D**). These studies show that RGS6 is co-localized in VTA DA synaptic terminals in the NAc and its loss leads to upregulated DAT expression which functions to extinguish mesolimbic neurotransmission.

## 4. Discussion

Alcohol is the most-commonly abused drug worldwide and AUDs impart a significant socioeconomic burden on the U.S. healthcare system, affecting ∼12% of the population. Chronic EtOH consumption promotes significant and progressive neuroadaptations within the mesolimbic circuit, often leading to dependence and abuse. AUDs are a chronic disorder with a high rate of relapse with 85% of patients treated for AUDs relapsing even after long periods of abstinence(39). While it is well known that the mesolimbic DA system mediates the behavioral responses to EtOH and other drugs of abuse(21), the mechanisms controlling VTA DA transmission and their impact on EtOH seeking behavior, dependence, and relapse are still not fully understood.

Here, we provide new evidence that RGS6 in VTA DA neurons plays a critical role in promoting EtOH consumption, preference, behavioral reward, and relapse. Collectively our findings strongly implicate RGS6 as an essential regulator of DA bioavailability in the mesolimbic circuit. Using a novel mouse model, we showed that loss of RGS6 from VTA DA neurons was sufficient to reduce EtOH consumption, preference, behavioral reward, and relapse in a manner indistinguishable from that seen in RGS6^−/−^ mice. Importantly, RGS6 was co-expressed with DAT in VTA dopaminergic terminals and RGS6^−/−^ mice showed significant upregulation of DAT, which functions to extinguish DA transmission. Consistent with these findings, we previously found that reduced EtOH consumption of RGS6^−/−^ mice was completely reversed by singular treatment with a DAT inhibitor, and that RGS6^−/−^ mice exhibit a reduction in striatal DA levels(32). Our findings suggest that RGS6’s negative regulation of GPCR-Gα_i/o_-DAT signaling promotes DA transmission contributing to EtOH consumption, behavioral reward, and relapse (**Fig. 5**). *RGS6* is an AUD gene in humans(33) and the current findings identify a locus of RGS6 action in the mesolimbic circuit.

**Figure 5.**
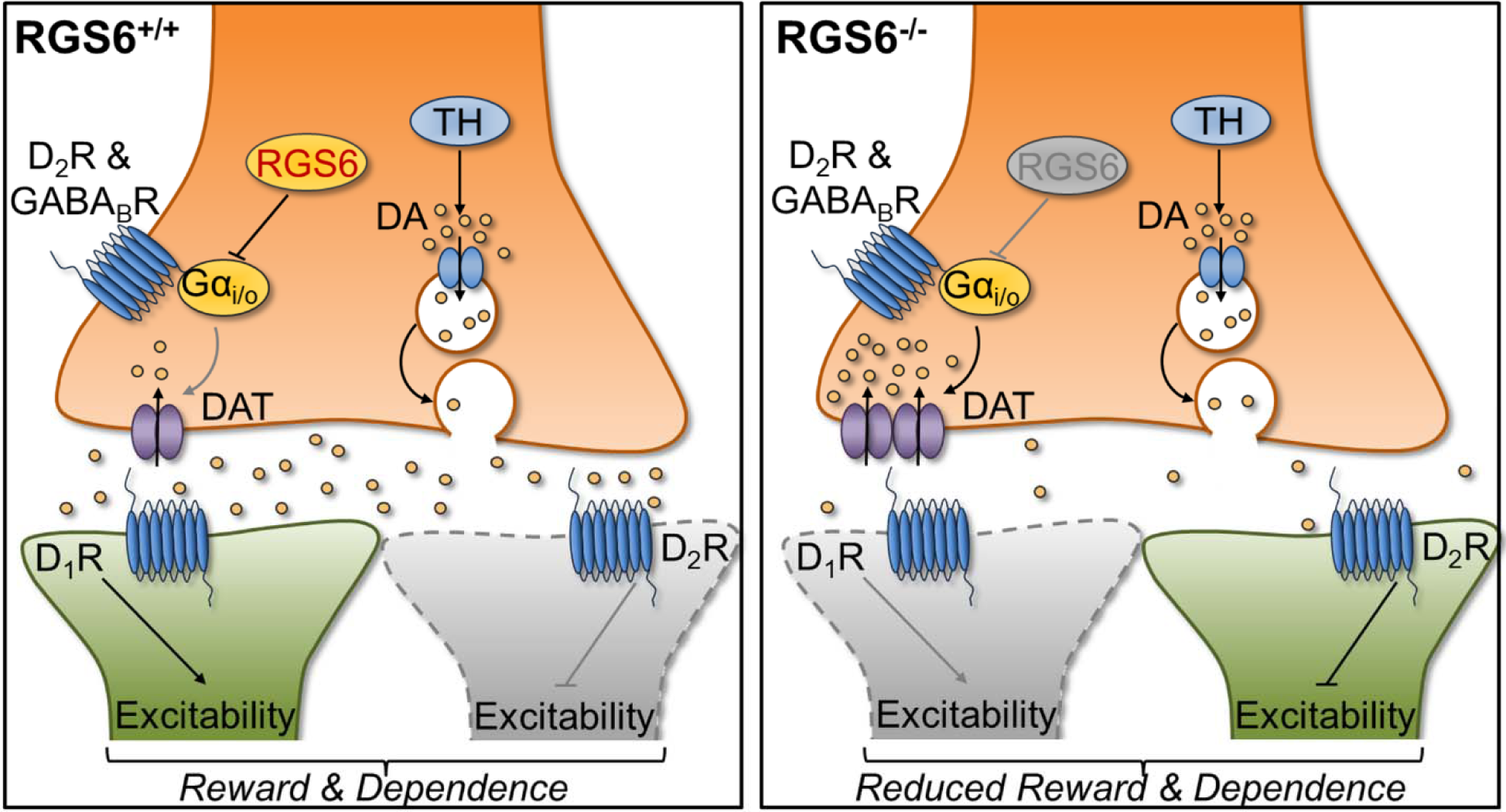
Proposed role of RGS6 in the mesolimbic circuit. By inhibiting GPCR-Gα_i/o_ signaling-mediated DAT upregulation/activation, RGS6 promotes EtOH consumption and reward behaviors (*left*). As a result, RGS6 deletion from VTA DA neurons leads to reduced EtOH consumption and reward by promoting DAT upregulation at synaptic terminals reducing the amount of synaptic DA release to the NAc (*right*).

We found that RGS6 is robustly and selectively expressed in DA neurons in the VTA of both humans and mice. Importantly, RGS6^−/−^ and RGS6 KO^VTA^ mice did not exhibit loss of VTA DA neurons. Thus, the decreased EtOH consumption, behavioral reward and relapse exhibited by these mice are not due to dopaminergic neurodegeneration. This is important, as we previously reported that RGS6^−/−^ mice display age-dependent degeneration of DA neurons in a neighboring midbrain region, the substantia nigra (SNc)(35), leading to Parkinson’s-like motor deficits(28). Thus, VTA dopaminergic degeneration is not responsible for the diminished EtOH consumption and reward behaviors seen in RGS6^−/−^ and RGS6 KO^VTA^ mice.

Our use of tamoxifen-inducible DAT-Cre mediated deletion of RGS6 from VTA DA neurons was based on numerous considerations. First, Lammel *et al.*(36) demonstrated overwhelming (96 +/− 1%) specificity of DAT-Cre expression in TH+ neurons in the midbrain and the relative lack of specificity of TH-Cre expression (48-59%) in TH+ neurons in this region. Second, VTA neurons projecting to the NAc have high DAT levels compared to those projecting to non-striatal sites. Indeed, we found that RGS6 was robustly co-expressed with DAT in synaptic terminals in the NAc. Third, inducible deletion of RGS6 allows for expression of RGS6 throughout development, avoiding any potential compensatory changes occurring due to embryonic deletion of RGS6. And finally, these mice provided a unique opportunity to investigate the role of RGS6 in VTA DA neurons to EtOH relapse behavior following extinguishment of behavioral reward.

Our findings provide new mechanistic insight into EtOH behavioral reward and relapse behavior by showing that RGS6 in VTA neurons is a critical mediator of these behaviors. Most patients with AUD experience relapse despite long periods of abstinence, with EtOH craving triggered by environmental cues and stress(41, 54, 55). Therefore, we employed two CPP-based extinguishment-reinstatement paradigms that model susceptibility to drug-induced reward and relapse (42)to study the role of RGS6 in these processes. We examined reinstatement of CPP with a single injection of EtOH following a withdrawal-like extinguishment phase. We performed RGS6 deletion from VTA DA neurons both before and after EtOH-induced CPP was established. Both RGS6^−/−^ mice and mice with knockout of RGS6 in VTA DA neurons before EtOH CPP studies were performed showed a significant and equivalent loss in both EtOH-induced CPP and its re-instatement. Strikingly, deletion of RGS6 from VTA DA neurons after EtOH-induced CPP was established led to a comparable (∼50%) loss of relapse behavior in these mice. Thus, RGS6 expression in VTA DA neurons regulates EtOH-induced reward and relapse susceptibility. Further our results show that the need for RGS6 in VTA DA neurons to establish EtOH reward is distinct from its role in promoting relapse. Because these two processes go hand-in-hand with patients with AUD and RGS6 deletion does not occur following establishment of EtOH reward and dependence, it seems likely that RGS6 is required for a common process for both behaviors, likely DA neurotransmission. It is possible that this common element is RGS6 effects on DAT synaptic expression, given that drugs that stimulate DA transmission like DAT inhibitors effectively re-instate CPP to cocaine(56). Finally, our results implicate VTA DA neurons in the process of EtOH-induced CPP relapse. Though we are unaware of previous studies implicating the VTA in reinstatement of EtOH CPP in mice, dopamine and mesolimbic structures have been found to play a role in reinstatement of CPP to other drugs of abuse(40, 42).

Though the present study is the first to examine effects of RGS6 loss in VTA DA neurons on EtOH seeking behavior, preference, reward, and relapse, DeBaker *et al.*(57) recently examined effects of AAV gRGS6 delivery to the VTA on D_2_R signaling in DA neurons and binge EtOH drinking in DATCre:Cas9 mice. Though deletion of RGS6 from VTA DA neurons was not confirmed, gRGS6-treated mice showed increased D_2_R-mediated somatodendritic currents in VTA DA neurons, while RGS6 overexpression reduced these currents. In addition, while treatment of these mice with gRGS6 had no effects on EtOH (20%) consumption during a 3day drinking in the dark paradigm (2hr access to EtOH), female but not male gRGS6-treated male mice exhibited an ∼25% reduction in “binge drinking” during 4hr access to EtOH on day 4. In contrast, a reduction in binge drinking in RGS6^−/−^ mice was observed in both males and females, as we found(32). Thus, this study reveals a unique sex-specific effect of RGS6 loss from VTA DA neurons on binge drinking and provides further support for the proposed function of RGS6 as a critical negative regulator of presynaptic D_2_R signaling in VTA DA neurons (**Fig. 5**), first suggested by our studies in mice with global RGS6 loss(32).

In summary, our studies have illuminated a critical role for RGS6 in the mesolimbic circuit in promoting EtOH seeking, preference, behavioral reward, and relapse. Our results have shown that the role of RGS6 in VTA DA neurons in promoting behavioral reward to EtOH is distinct from its role in promoting relapse. We propose that the common element in these behaviors may be RGS6 modulation of VTA DA transmission, with RGS6 functioning to negatively regulate GPCR control of a Gαi/o-DAT pathway (**Fig. 5**). Indeed, RGS6 is co-expressed with DAT in VTA synaptic terminals and RGS6 loss leads to upregulated DAT expression. These findings are entirely consistent with our observations that diminished EtOH seeking in RGS6^−/−^ mice is associated with diminished striatal DA content, partly reversed by antagonism of GABA_B_Rs or D_2_Rs, presynaptic GPCRs in VTA neurons whose activation diminishes EtOH consumption(14, 21, 58), and completely reversed by DAT inhibition(32). Indeed, RGS6 loss from VTA DA neurons recapitulates the behavioral effects we observed in RGS6^−/−^ mice. Thus, these studies identify the mesolimbic pathway, the brains major reward pathway, as the locus of RGS6 action in modulating EtOH behaviors. Our studies identify RGS6 as a potential therapeutic target for behavioral reward and relapse to EtOH. Given that deletion of RGS6 from VTA DA neurons after established behavioral reward significantly prevented reinstatement induced relapse, RGS6 inhibition could significantly prevent relapse. These findings provide new mechanistic insights that are consistent with the identification of RGS6 as an AUD gene in humans.

## Author Contributions

M.M.S., M.A.W., N.S.N., and R.A.F contributed to preparation of the manuscript (writing/editing/figure compilation) and experimental design. M.M.S. executed all behavioral experiments and statistical analyses. M.M.S. and Z.L. performed histology. M.M.S. and M.A.W. designed all behavioral studies. M.M.S. and J.Y designed and created mouse strains used for all studies and performed immunoblotting quantification.

## Supplementary Material

### Methods

All mice were housed in a room maintained at 22^°^C and 20-30% humidity on a reverse light cycle (12hrs light: 12hrs dark-lights off at 8:00am) and were given *ad libitum* access to food and water. Mice for behavioral studies were individually housed starting 1 week prior to behavioral assessments. All *in vivo* experiments were approved and performed in accordance with guidelines set by the Institutional Animal Care and Use Committee (IACUC) at the University of Iowa.

#### 2.2 Behavioral Studies

All behavior analyses were conducted during the dark phase of the light cycle and performed in rooms as dimly lit as possible to avoid interrupting the sleep cycle and normal behavior patterns of mice. All mice were handled for 5 consecutive days prior to starting behavioral studies, and mice were handled in the respective behavior room 24hrs prior to beginning trials. On each day of training/testing, mice were acclimated to the room for 60min prior to beginning the trial and the experimenter remained out of sight for all behavioral trials.

##### 2.2.1 Short-term EtOH consumption and preference

Short-term, two-bottle choice (2BC) EtOH consumption and preference were measured in RGS6^+/+^, RGS6 WT^VTA^, RGS6 KO^VTA^, and RGS6^−/−^ mice, as we previously described(32). Mice were individually housed 5 days prior to beginning 2BC studies. For the first three days of 2BC, mice received two bottles of tap water. On day 4, mice received one bottle of tap water and one bottle of 8% (vol/vol) EtOH for two days. Finally, mice received two bottles of 8% (vol/vol) EtOH for the final 2 days (**Fig. 3A**). Bottle volumes (mL) and mouse weights (g) were recorded daily and bottle positions were alternated daily to avoid place preference bias. EtOH consumption was normalized to individual mouse body weights.

##### 2.2.2 Conditioned Place Preference (CPP), Extinguishment, and Reinstatement

CPP was performed as we previously described(32). Prior to conditioning, mice were habituated to the chambers of the CPP box by injecting saline (i.p.) and placing them in the center of the box, recording their exploration and side preferences for 15 min. Mice were subsequently conditioned (12 trials total, 5 min/trial, total of 6 trials per treatment/chamber pairing) to associate EtOH (2 g/kg i.p.) with their non-preferred side, and saline with the opposite side. A subset of animals that didn’t undergo EtOH conditioning (*i.e.* saline only) were included as controls. 24hrs following conditioning, mice received singular saline injections (i.p.), were placed in the center chamber, and allowed to freely explore both chambers for 15min. Their time spent in the EtOH chamber side was recorded.

Following CPP, mice underwent an extinguishment paradigm where they received single saline injections (i.p.) and were allowed to freely explore both chambers for 15min. Mice were tested daily until they displayed no preference to the EtOH chamber (≤ 450 sec). After extinguishment had been established, mice were given a single injection (i.p.) of EtOH (2 g/kg), placed in the center of the chamber, and allowed to freely explore both chambers for 15min (reinstatement phase).

#### 2.3 Immunofluorescence (IF)

In preparation for IF, mouse tissues mice were perfused as we previously described(28, 59). Briefly, mice were first euthanized with isoflurane and immediately perfused (cardiac) with ice-cold 1X PBS followed by 4% paraformaldehyde (PFA). Brain tissues were removed, postfixed in 4% PFA at 4^°^C overnight, and finally sunk in a 30% sucrose solution (in 1X PBS). 10 µm coronal DG sections were collected (Olympus cryostat) and mounted. Tissue sections were blocked for 1hr at 4^°^C in 1X PBS + 10% goat serum + 0.5% Triton X-100 (blocking solution). After blocking, tissue sections were incubated overnight at 4^°^C in rabbit-anti-RGS6 (homemade polyclonal antibody against the entire RGS6 protein(26, 30, 32, 60), 1:100, mouse-anti-TH (1:100, Millipore, MAB318), or mouse-anti-DAT (1:100, Millipore MAB669) primary antibody diluted in the blocking solution. Following overnight incubation, tissues were washed 5 times at room temperature in 1X PBS and subsequently incubated for 1hr at 4^°^C in Alexa Fluor 488/568/647-conjugated secondary fluorescent antibodies. Slices were mounted with ProLong Diamond Antifade Mountant with DAPI (Fisher Scientific) and imaged using an Olympus FV3000 confocal microscope. Cell counts were performed as previously described using ImageJ (35).

#### 2.4 Western Blotting (WB)

Mouse VTA and NAc tissue samples were lysed using 100µL of RIPA buffer (150mM NaCl, 1% NP-40, 0.5% sodium deoxycholate, 0.1% sodium dodecyl sulphate, 50mM Tris-HCl: pH 8) containing protease (Roche) and phosphatase (Sigma) inhibitors. Lysed tissues + RIPA buffer were placed in pre-chilled tubes, vortexed (∼30 sec), and allowed to incubate on ice (5 min). Following incubation, lysates were spun at 8000rpm in a 4LC centrifuge (10 min). The supernatant was subsequently transferred and divided between empty (pre-chilled) tubes for protein quantification (BioRad DC Protein Assay, Cat# 500-0116) and tubes containing 4x Loading Buffer for WB analysis. Samples in loading buffer were boiled (5 min) prior to loading into 10% Mini-PROTEAN® TGX Stain-Free Precast Gels (BioRad, Cat# 4568036). Western blots were probed with a home-made rabbit anti-human RGS6 primary antibody (1:10,000) against the whole protein(26, 60), as well as monoclonal DAT (1:1000, Millipore, MAB669), primary antibody. LI-COR Odyssey goat anti-mouse (Cat# 926-32220 and 926-32210) and goat anti-rabbit (Cat# 926-32211 and 926-68071) were used at a 1:5000 dilution for protein visualization with the Odyssey system. Quantification of relative DAT expression was performed as previously described (60).

### Supplementary Figure Legends & Figures

**Supplemental Figure 1.**
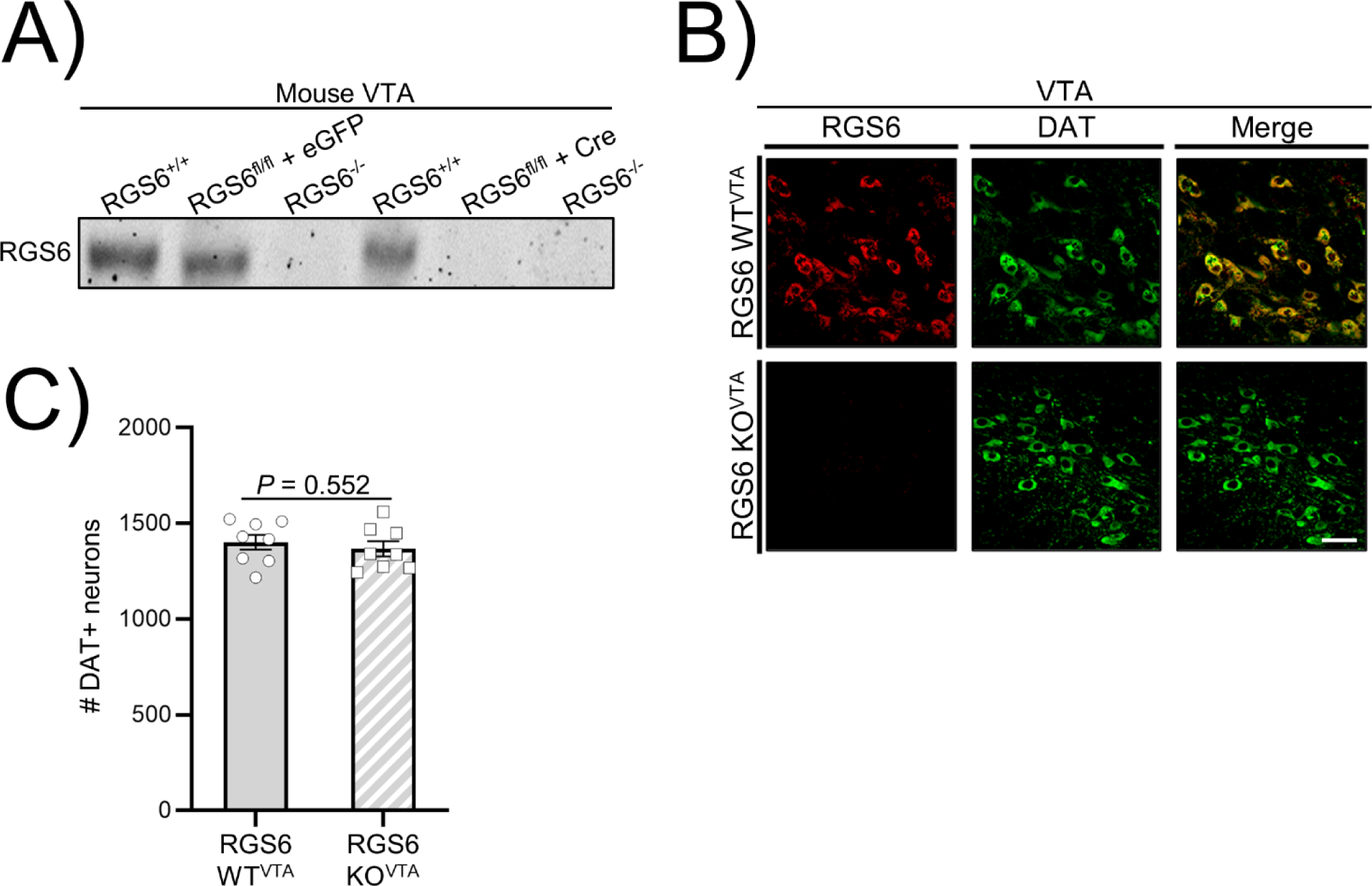
Inducible RGS6 deletion from VTA dopamine neurons in mice. **A)** Western blot showing RGS6 deletion from the VTA of RGS6^fl/fl^ mice by stereotactic administration of AAV-Cre-eGFP but not AAV-eGFP (left panel) into the VTA. RGS6 expression in RGS6^+/+^ and RGS6^−/−^ mice serve as controls. **B-C)** Immunofluorescent (IF) analysis (**B**) and quantification (**C**) of tamoxifen-mediated deletion of RGS6 (red) from TH^+^ DA neurons (green) in the VTA of 3mo. RGS6^fl/fl^;DATiCreER mice. Scale bar represents 40 μm.

**Supplemental Figure 2.**
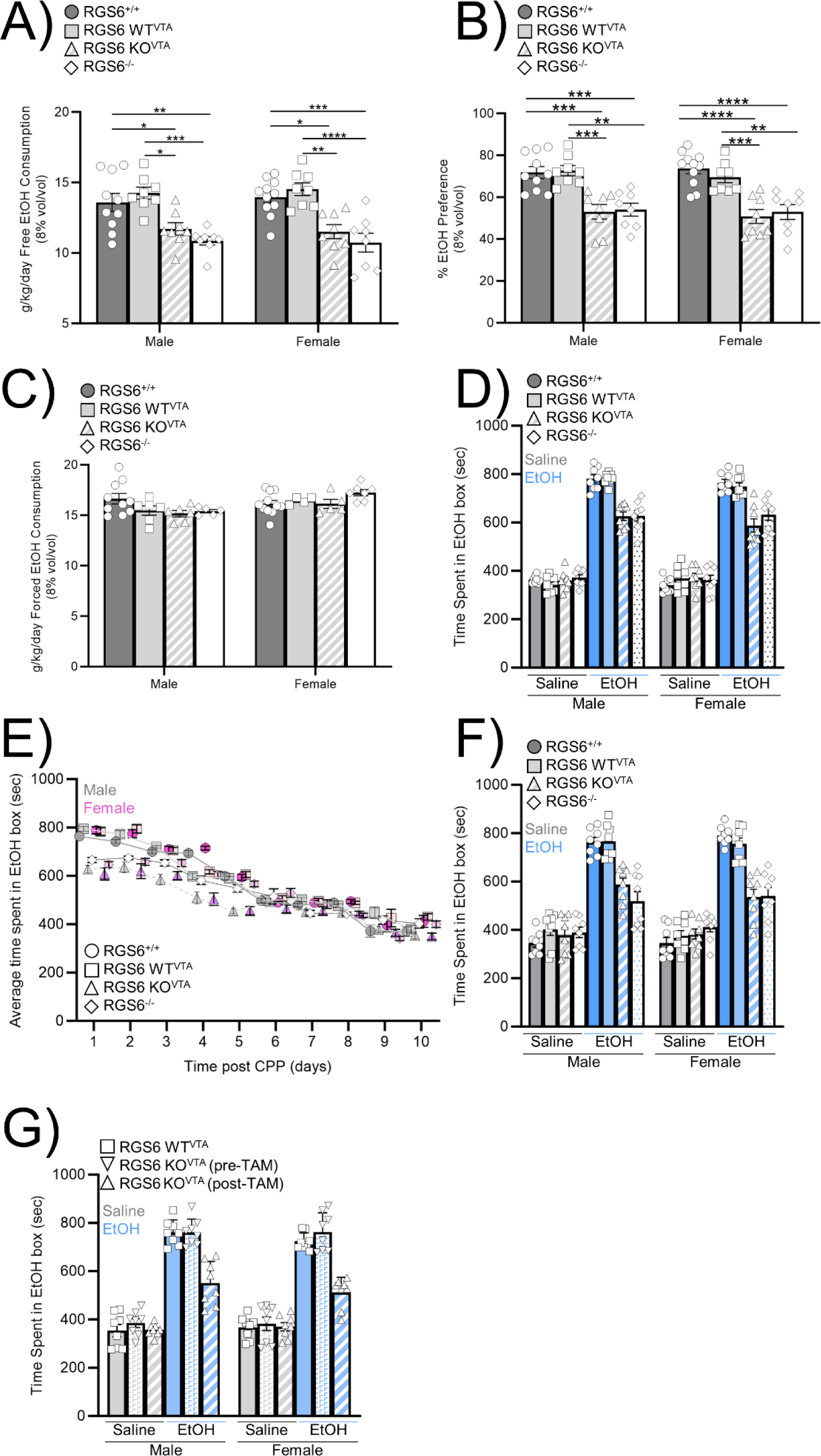
VTA loss of RGS6 in VTA DA neurons diminishes EtOH drinking, reward, and relapse behaviors in male and female mice. **A-G)** Data here show data from **Figs. 3** and **4** organized by male and female mice for short term EtOH consumption and preference (**A-B**), forced consumption (**C**), CPP (**D**), extinguishment (**E**), and reinstatement in mice with RGS6 deletion before **(F)** and after **(G)** established CPP. No significant effect of sex or its interactions were observed across all behaviors. \**P* ≤ 0.05, \*\**P* ≤ 0.01, \*\*\**P* ≤ 0.001, \*\*\*\**P* ≤ 0.0001.

